# Sourmash accurately profiles strains from long-read metagenomes preserving gene content

**DOI:** 10.64898/2026.11.03.763925

**Authors:** Lune Angevin, Josipa Lipovac, Pierre Peterlongo, Emeline Roux, Riccardo Vicedomini

## Abstract

**Background:** Metagenomic strain-level profiling remains challenging because closely related bacterial strains often share more than 99% average nucleotide identity (ANI). However, they can differ substantially in gene content and functional potential. Sourmash uses k-mer sketches with FracMinHash to compare sequencing reads against reference databases, which makes it an interesting approach for strain-level profiling. Because it compares sketches rather than full alignments, it enables fast searches against large reference databases. However, its ability to recover biologically relevant strains and preserve community gene content has not been specifically assessed. Here, we evaluated the performance of sourmash for strain-level identification abundance, functional gene content recovery, and abundance estimations using five mock communities of increasing complexity with known composition.

**Results:** We first assessed whether nucleotide similarity reflects functional similarity between strains. Gene content similarity was generally consistent with ANI but revealed important divergences between closely related strains. For example, *Escherichia coli* strains sharing more than 98% ANI showed gene content similarities ranging from 20% to 57%, highlighting that nucleotide similarity alone is insufficient to assess functional proximity and consequently the accuracy of strain identification. Above sufficient genome coverage, sourmash identified strains either exactly or with closely related strains sharing highly conserved gene content. These closely related strains provided a biologically relevant representation of the expected functional potential of the mock communities. Gene content recovery was highly accurate, with F1 scores ranging from 0.942 to 0.998 across the five mock communities. Genome abundances estimated against sourmash-identified genomes closely matched the expected community composition: Pearson correlation coefficients ranged from 0.823 to 1 at strain level and from 0.978 to 1 at species level.

**Conclusions:** Our results show that sourmash provides a biologically relevant approach for strain-level characterisation in microbial ecosystems. Even when the exact strain could not be identified, closely related strains retained most of the expected gene content and abundance estimations were accurate, supporting the use of sourmash for downstream functional profiling or metabolic network reconstruction.

## Background

Metagenomics is a crucial approach for studying microbial ecosystems such as the gut microbiota. The development of high-throughput sequencing has particularly contributed to the progress of this field (1). For many years, 16S rRNA gene sequencing has been the standard approach. However, because it is based on a small fraction of the bacterial genome (roughly 500 bp per genome), its taxonomic resolution is limited at the genus level. On the contrary, shotgun metagenomic sequencing is an untargeted approach, which captures DNA from across the genomes of microorganisms present in the sample. With sufficient depth, it can provide access to single-nucleotide variants and structural variants, enabling potential identification at the bacterial strain level (2).

Genetically, microbial species are defined by an average nucleotide identity (ANI) (3) of 95-97%, while conspecific strains (*i.e.*, strains belonging to the same species) typically share ANI values above 99% and up to 99.9% (2,4). Therefore, two strains share a large proportion of their genomes but differ in small genetic variations. Identifying bacteria down to the strain level is particularly important in a health context, as two strains can vary from beneficial or commensal to pathogenic, as it has been demonstrated for *Escherichia coli*, *Listeria monocytogenes*, and *Propionibacterium acnes* (2,4–6). However, the similarity between two strains depends not only on their nucleotide sequence similarity. Nucleotide variations can lead to differences in gene content and consequently to functional differences. For downstream applications based on functional profiling, such as metabolic network reconstruction, an essential question is whether a closely related strain remains a biologically meaningful proxy for the expected strain by preserving its gene content. The genes collection of a genome determines its functional potential, defined here as the biological functions that can potentially be encoded by its genes.

The choice of the sequencing technology affects the ability to access strain-level identification. Short reads, ranging from a few dozen to a few hundred bases, are more advantageous in terms of cost and accuracy, but they can not capture large structural variants or complex regions such as repeats (2,7). Long reads, on the other hand, vary from thousands to hundreds of thousands of bases. Modern long-read sequencing technologies, such as PacBio HiFi and Oxford Nanopore R10, have reduced error rates below 1% (8). These improvements are driving the increasing adoption of long-read metagenomics. However, as long-read metagenomic datasets continue to grow, scalable approaches leveraging these technologies are needed to efficiently identify strains from microbial communities.

Beyond sequencing, the characterisation of metagenomic datasets represents a computational challenge at the strain level. Several strategies have been developed for strain-level profiling. Marker-gene- and SNV-based methods, such as StrainPhlAn (9) and inStrain (10), characterise strain diversity by analysing nucleotide variants. While these approaches provide high-resolution strain comparisons, they are based on accurate alignments and primarily designed for variant analysis rather than whole-genome similarity. In a similar way, reference-based read assignment methods, such as MORA (11) or StrainRefine (12), accurately identify strains but require a lot of memory. For example, MORA may require ∼1TB of RAM to run on a dataset of 5M reads (13). *De novo* assembly methods can provide an alternative but may require high-quality reads and sufficient coverage. A recent workflow, MADRe (13), combines *de novo* assembly and a contig-to-reference mapping to create a reduced database, before using a probabilistic read mapping reassignment. Even if it shows good results in strain identification, it produces up to 75% of false positives (12). Furthermore, as metagenomic datasets and reference databases are always increasing in size, scalable approaches able to compare large numbers of genomes are required. The use of k-mers (subsequences of length k) may provide solutions to the issue of running time. The representation of reads as a set of k-mers enables much faster comparisons between genomes. For instance, Mash (14), based on k-mer sketches, was used to sketch and compare 54k genomes from RefSeq (1.5 billion pairwise distances computed) in 33 CPU hours. This achievement would be infeasible even with the fastest tools computing exact distances. Extrapolating from Edlib (15) results, for example, would require hundreds of centuries of CPU time. Representing reads as k-mers is implemented in ORI (16), a method conceived to identify strain-level bacteria, yet not designed for complex metagenomes. Moreover, taking into account all the k-mers of a large dataset can lead to storage limitations (2,7). To reduce both storage and running time, sketching algorithms such as MinHash have been developed (14). They enable the subsampling of k-mers while preserving the initial similarity between the datasets (17).

Sourmash (17), using sketching with FracMinHash (18), allows a scalable comparison of metagenomic datasets with large reference genome databases. While it was initially developed for large-scale sequence comparison, its performance for strain-level identification and recovery of gene content has not been specifically evaluated. Here, we assessed the ability of sourmash to identify bacterial strains present in metagenomic samples from the human gut, a well-characterised ecosystem, using a comprehensive reference database. All reference-based approaches are limited by the existence of representative genomes, as strains actually present in a sample may be missing in the databases. Consequently, when the exact strain is absent from the database, identifying closely related genomes preserving gene content remains an important challenge. Hence, beyond exact strain identification, our goal was to determine if the identified strains preserved the gene content of the target strains. Therefore, we assessed the accuracy of strain identification and also the similarity of gene content. Furthermore, we assessed whether sourmash-identified genomes allowed to preserve abundances. These tests were carried out using mock communities, composed of human gut or oral bacteria, in order to take advantage of ground truth information.

## Methods

### Datasets

Mock communities, artificial mixtures of microorganisms in known and controlled proportions, are useful for assessing the quality of an analysis pipeline. In this study, five mock communities of increasing complexity were selected. The ZymoBIOMICS HMW community contains seven bacterial strains from seven different species of equal abundance (14%) and one yeast (2%). The ZymoBIOMICS ORAL community contains 12 bacterial strains from 12 different species, with abundances ranging from 1% to 16%. The ZymoBIOMICS GUT community contains 18 bacterial strains from 13 different species, including five strains of *Escherichia coli*, as well as two yeasts and one archaea. Abundances range from 0.0001% to 14%, and the five *Escherichia coli* strains have the same abundance of 2.8%. The Nanopore R10.4 sequencing long reads for these three communities were retrieved from the MicroBench repository (https://github.com/Kirk3gaard/MicroBench). The CONCAT community was generated as described in Shaw *et al.* (19). It was built by concatenating the sequencing datasets of the three previous communities (*i.e.*, HMW, ORAL, and GUT) with those of 14 isolated strains, for a total of 49 bacterial strains from 39 different species, two yeasts, and one archaeon. Datasets for the 14 isolated strains were downsampled to 10% of their original size, unless the total data was already below 1 Gb. After concatenation, the resulting dataset was further downsampled to 25% of its size, yielding the CONCAT community used in this study. The last dataset, PANTAX, is a simulated community from Zhang *et al.* (20) and, as opposed to the previous ones, its reads were generated using CAMISIM (21) with the Nanopore R10.4 sequencing model. It represents 60 bacterial strains from 30 different species, and their abundances range from 0.30% to 6.43%. Detailed composition of each mock, including strains accessions number, can be found in Supplementary Data 1. ANI scores were calculated between all strains of each mock community using fastANI (3).

### Reference genome databases

Two databases of different sizes were evaluated. The largest one, RefSeq-CG, comprises all the RefSeq complete bacterial genomes. The genomes were downloaded from the NCBI using Kraken2 (22) with the command *kraken2-build* with the option *--download-library bacteria*. It built a database of 248 GB, containing 144,488 reference sequences. This database was created in November 2025.

The other database, MADRe-RD, is a reduced version of the RefSeq-CG database generated using the tool MADRe (13) independently for each mock community. MADRe selects a subset of the genomes of the input database based on an assembly of the sequencing reads and contig-to-reference mapping. MADRe was run with the parameter *--reads_flag ont* as the input consisted of Nanopore reads. Depending on the mock community, the reduced databases ranged from 55 MB with 27 reference sequences to 1.8 GB with 480 reference sequences (Table 1). This approach was evaluated to assess whether a reduced database could serve as a pre-filter to improve runtime and precision.

**Table 1:** Characteristics of the five mock communities (Nanopore R10.4 reads).

| **Mocks** | **HMW** | **ORAL** | **GUT** | **CONCAT** | **PANTAX** |
|---|---|---|---|---|---|
| **Read type** | Sequenced |  |  |  | Simulated |
| **Number of bacterial species** | 7 | 12 | 13 | 39 | 30 |
| **Number of bacterial strains** | 7 | 12 | 18 | 49 | 60 |
| **Smallest coverage** | 265X | 9X | 0.01X | 0.01X | 0.97X |
| **Highest coverage** | 1164X | 711X | 4090X | 1021X | 20X |
| **Number of bases** | 20.19 Gbp | 7.53 Gbp | 46.50 Gbp | 20.56 Gbp | 1.27 Gbp |

**Table 2:** Characteristics of reference genome databases.

| **Mocks** | **HMW** | **ORAL** | **GUT** | **CONCAT** | **PANTAX** |
|---|---|---|---|---|---|
| **RefSeq-CG database size** | 248 GB |  |  |  |  |
| **RefSeq-CG database number of sequences** | 144,488 |  |  |  |  |
| **MADRe-RD database size** | 153 MB | 55 MB | 641 MB | 1.8 GB | 469 MB |
| **MADRe-RD database number of sequences** | 47 | 27 | 158 | 480 | 163 |
| **Proportion of mock bacterial strains present in MADRe-RD** | 4/7 | 9/12 | 13/18 | 24/49 | 49/60 |

### Sourmash-based analysis

We evaluated sourmash v4.9.4 (17) on strain identification in human gut microbiota metagenomes. As shown in Figure 1, the workflow consisted of three steps: reference database sketching, samples sketching, and similarity computation. We added a step to filter the list of genomes identified with sourmash.

**Figure 1:**
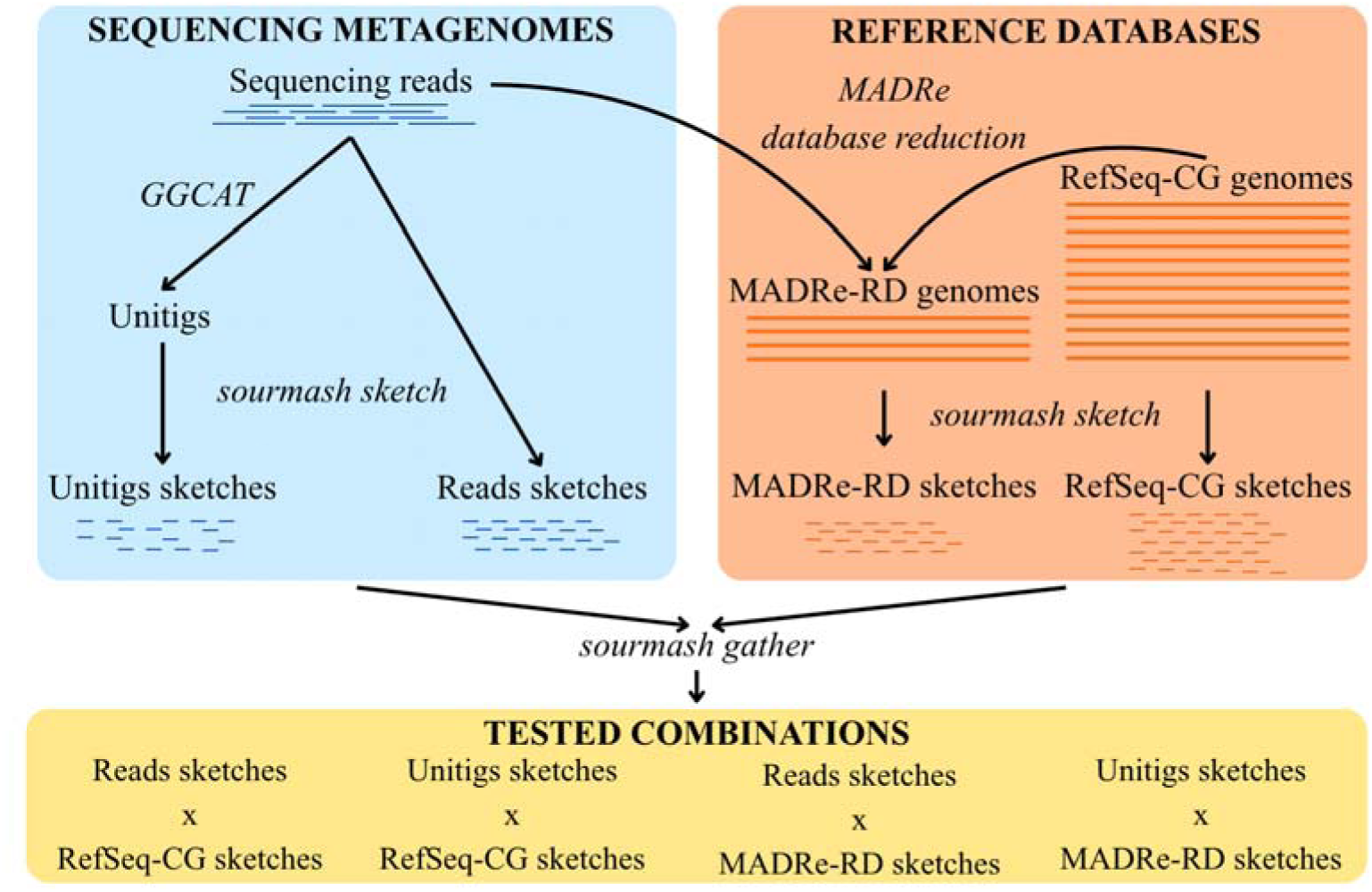
Overview of the four configurations to evaluate sourmash using read- or unitig-based sketches and RefSeq-CG or MADRe-RD database sketches. The sequencing metagenomes on the left panel refer to the data to characterise.

#### 1. Database sketching

The RefSeq-CG database and the five MADRe-RD databases were transformed into k-mers sketches with MinHash using the command *sourmash sketch dna*. The option *--singleton* was used, which allows creating signatures for each sequence. The sketches were then indexed using RocksDB (23).

#### 2. Read and unitig sketching

For each mock community, we considered either the raw sequenced reads or k-mers seen at least twice, and represented as unitigs. Since sequencing errors introduce erroneous low-abundance k-mers, building unitigs removes such k-mers, reduces data redundancy, and lowers the computational resources required. Unitigs were generated with GGCAT v2.0.0 (23) with default parameters and k set to 31, which discards unique k-mers. Sketches were generated using the command *sourmash sketch dna* with the option *-p k=31,scaled=1000,abund* for sequencing reads, and *-p k=31,scaled=1000* for unitigs.

#### 3. Similarity search

The command *sourmash gather* was used to identify the reference genomes of the database sharing k-mers with sample sketches. In the case of unitigs, the option *--ignore-abundance* was used because the sketches were generated without the *abund* parameter and so did not contain k-mer abundance information.

#### 4. Filtering the results

Sourmash-identified genomes were filtered on the *f_match_orig* metric, defined as the fraction of query k-mers matching those of the reference genomes. For simplicity, this value is referred to as the “*reference fraction*” throughout this manuscript. Only strain genomes with a *reference fraction* above 0.5 were retained, meaning that at least 50% of their reference genome is estimated to be present in the metagenomic data. Several thresholds between 0.5 and 1 were tested, as detailed in the Results and Discussion section.

### Gene content analysis

In order to compare strains in terms of gene content, the proteomes of reference genomes proposed by sourmash and of mock community genomes were retrieved from NCBI. To have homogeneous annotations, all proteomes were reannotated with EggNOG-mapper v2.0.1 (24). The diamond mode was used with the bacterial EggNOG database. The other parameters were *--seed_ortholog_evalue 0.001 --seed_ortholog_score 60 --query_cover 20*. For each proteome, EggNOG orthologous groups were extracted from the annotation file.

### Performance metrics for assignation and abundances

Throughout this study, *exact strain* identification refers to cases where sourmash identifies the same reference genome as the one present in the mock community. If this is not the case, we define the “*closely related*” strain of an expected strain as the one predicted by sourmash whose gene content is closest to that of the expected strain.

Several metrics were calculated to assess sourmash strain identification results. Gene content similarity was evaluated at two levels: between individual strain pairs, and globally between genome sets. For pairwise comparisons, we used the Jaccard similarity index of gene content between two genomes, computed as the intersection of their EggNOG orthologous groups divided by their union. For genome-set comparisons, the orthologous groups of strains identified by sourmash were compared with those of the expected strains in each mock community, and predictions were classified as true positives (TP), false positives (FP), and false negatives (FN). Recall is defined as the proportion of expected orthologous groups correctly identified. Precision is defined as the proportion of identified orthologous groups corresponding to expected orthologous groups. F1-score is the harmonic mean of precision and recall.

Coverages and abundances of each strain in the mock were calculated to establish the ground truth, as sequencing biases can affect the initial abundances provided by the mock designs (25). Reads were aligned to mock reference genomes with Minimap2 (26) to estimate genome coverage and only primary mappings were kept to avoid counting multiple alignments for the same read. Genome coverage was calculated as the sum of read lengths aligned to a genome divided by the genome length. Then, relative abundance was calculated as the coverage divided by the sum of coverages. Coverages and abundances of the genomes identified by sourmash were calculated the same way. To compare the ground truth abundances and the abundances of the genomes identified by sourmash, Pearson correlation coefficients were computed using the cor() function in R v4.4.3. It was calculated at the strain-level, using only exact strains identified by sourmash. It was also calculated at the species-level, by adding together the abundances of each strain of the same species.

## Results and Discussion

### Gene content similarity is consistent with ANI scores but reveals divergences among closely related strains

To study the link between nucleotide similarity and gene content similarity, we first characterised the genomes composing the mock communities. The simplest mock communities HMW and ORAL only contain one strain per species. In these two mock communities, almost all strains share an ANI score below 70% (Supplementary Figures 1A and 2A). We cannot provide precise values as the exact score is not calculated by fastANI below this threshold. In HMW mock community, *Escherichia coli* and *Salmonella enterica* have an ANI score of 82.24%, and in ORAL mock community, a Streptococcus cluster of *Streptococcus mitis*, *Streptococcus parasanguinis*, and *Streptococcus salivarius* has ANI scores ranging from 80.12 to 82.94%. ANI scores and gene content similarity are consistent (Supplementary Figures 1 and 2). Reflecting their low nucleotide sequence similarity, all strains in the HMW and ORAL communities share no gene content with one another (Supplementary Figures 1B and 2B).

The relationship between nucleotide similarity and gene content similarity becomes more informative in communities containing multiple strains of the same species. As a reminder, microbial species have an average nucleotide identity (ANI) of 95 to 97%, while conspecific strains can reach up to 99.9% identity (2,4). In the mock community GUT, the five *E. coli* strains share high ANI scores, ranging from 98.3% to 99.51% (Supplementary Figure 3A), highlighting the difficulty of distinguishing the closely related strains. However, high nucleotide sequence similarity does not necessarily imply high gene content similarity, as relatively small genomic differences can result in substantial changes in gene content through gene gain or loss. The *E. coli* strains have gene content similarities ranging from 20% to 57% (Supplementary Figure 3B).

The CONCAT community combines strains from HMW, ORAL, GUT, along with 14 isolated strains, resulting in 49 strains from 39 species. PANTAX, instead, contains two strains per species, for a total of 60 strains from 30 species. Being the two most complex mock communities, subsequent analyses focus on these two. Furthermore, among the mock communities based on real sequencing reads, the CONCAT provides the closest approximation to the diversity and strain complexity found in a real gut microbiota, even if it remains a simplified model. As expected, as far as the CONCAT community is concerned, strains belonging to the same species, such as *Escherichia coli*, *Enterococcus faecalis*, *Fusobacterium nucleatum*, L*isteria monocytogenes*, *Salmonella enterica*, and *Staphylococcus aureus* share high ANI values ranging from 91.2% to 99.9% (Figure 2A). All other strains share ANI scores below 70%. The observed patterns of gene content similarity are consistent with the ANI scores, with the highest gene content similarities found between strains of the same species (Figure 2B). For example, *Salmonella enterica* strains have a gene content similarity of 95%. Nevertheless, as in the mock community GUT, a high ANI score is not always linked to a high gene content similarity. The two strains of *Staphylococcus aureus* share an ANI value of 98.71% but only 22% of their orthologous groups. These observations highlight that nucleotide similarity provided by the ANI metric alone is insufficient for strain characterisation. For clinically important bacteria such as *E. coli* (2,4) and *L. monocytogenes* (5), differences in gene content can support important functional or pathogenic differences despite highly similar genomes. Similar observations can be made for PANTAX: strains of the same species share ANI scores over 95%, while strains of the same genus can reach ANI scores over 90%, as for the genus *Borrelia* (Supplementary Figure 4A). Looking at gene content similarity, no strain couple exceeds a gene content similarity of 49% (Supplementary Figure 4B).

**Figure 2:**
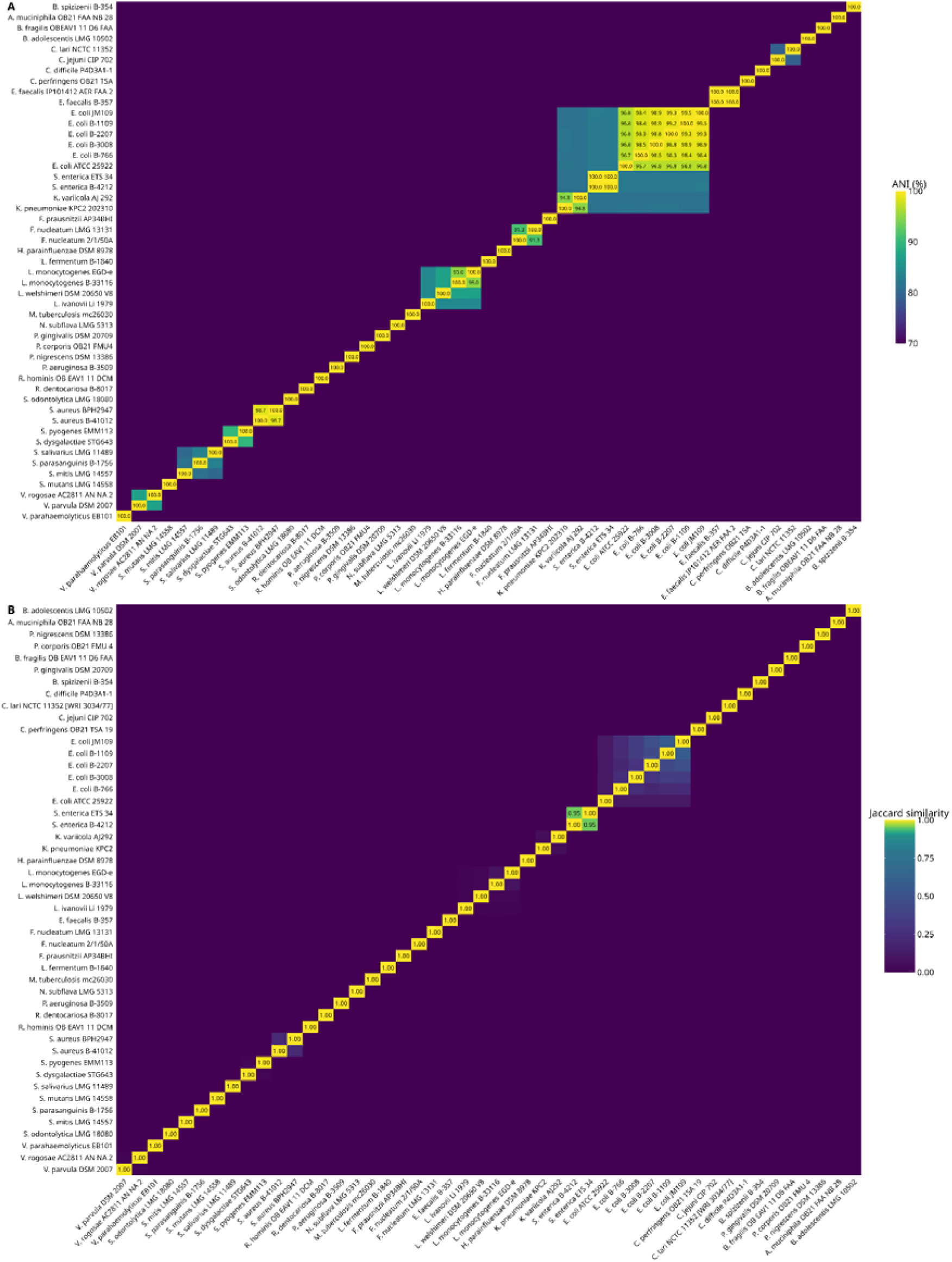
Characterisation of CONCAT mock community composition. **(A)** ANI scores heatmap of the strains of the mock community CONCAT. **(B)** Gene content similarity heatmap of gene content between CONCAT mock community strains. Strains of the same species have high ANI scores, over 90%. Below an ANI score of 70%, the exact score was not calculated by fastANI.

Taken together, these results confirm that closely related strains can differ substantially in their accessory gene content, despite a high genomic similarity. Therefore, gene content similarity provides complementary information to nucleotidic similarity and is thus relevant for evaluating strain-level identifications.

### Beyond exact strain identification: sourmash retrieves strains with conserved gene content

Sourmash was applied to all mock communities using the four possible combinations of reference databases and input data: read sketches or unitig sketches, combined with RefSeq-CG or MADRe-RD sketches (Supplementary Figure 5). Since unitigs were computed to preserve the k-mer content of the reads while discarding low-abundance erroneous k-mers, they were expected to improve results. However, unitig-based sketches did not outperform read-based sketches, likely due to the loss of low-abundance strain signal. The combination of sequencing reads and the RefSeq-CG database was retained for the analyses presented below, as it provided the best compromise between recall and precision (Supplementary Figure 5). For each strain identified by sourmash, gene content similarity was assessed by comparing its EggNOG orthologous groups with those of the expected strains in the corresponding mock community. This approach allows not only to evaluate strain identification accuracy, but also the biological relevance of closely related strains.

The ability of sourmash to identify strains, relying on k-mer sketching to compare genomes, is influenced by genome coverage in metagenomic datasets. In the CONCAT mock community, when coverage was below 8X, sourmash was unable to recover strains, either as exact or close, and some expected strains were not detected at all (Figure 3). Indeed, a sufficient number of reads is needed to get enough k-mers to generate a representative sketch and distinguish closely related genomes. Unlike alignment-based approaches, which require mapping of reads against references, k-mer sketching approaches are based on global similarity, which enables fast similarity searches. Therefore, they can rapidly identify related genomes even when the exact reference is not in the database. Above this coverage threshold, all strains of the mock were identified, either as the exact strain or as a closely related one. A gene content similarity of 1 indicates that an exact strain has been identified, while lower values reflect the identification of a closely related one. For the CONCAT mock community, those closely related strains have a gene content similarity of at least 0.89 (Figure 3). However, this non-exact similarity is not negligible. For example, two genomes containing 2000 genes and sharing 90% of their gene content may still differ by approximately 200 orthologous groups that are specific to each strain, which can represent important functional differences. Therefore, sourmash does not recover the exact functional potential of a strain but it still provides a relevant approximation of its gene content and functional potential.

**Figure 3:**
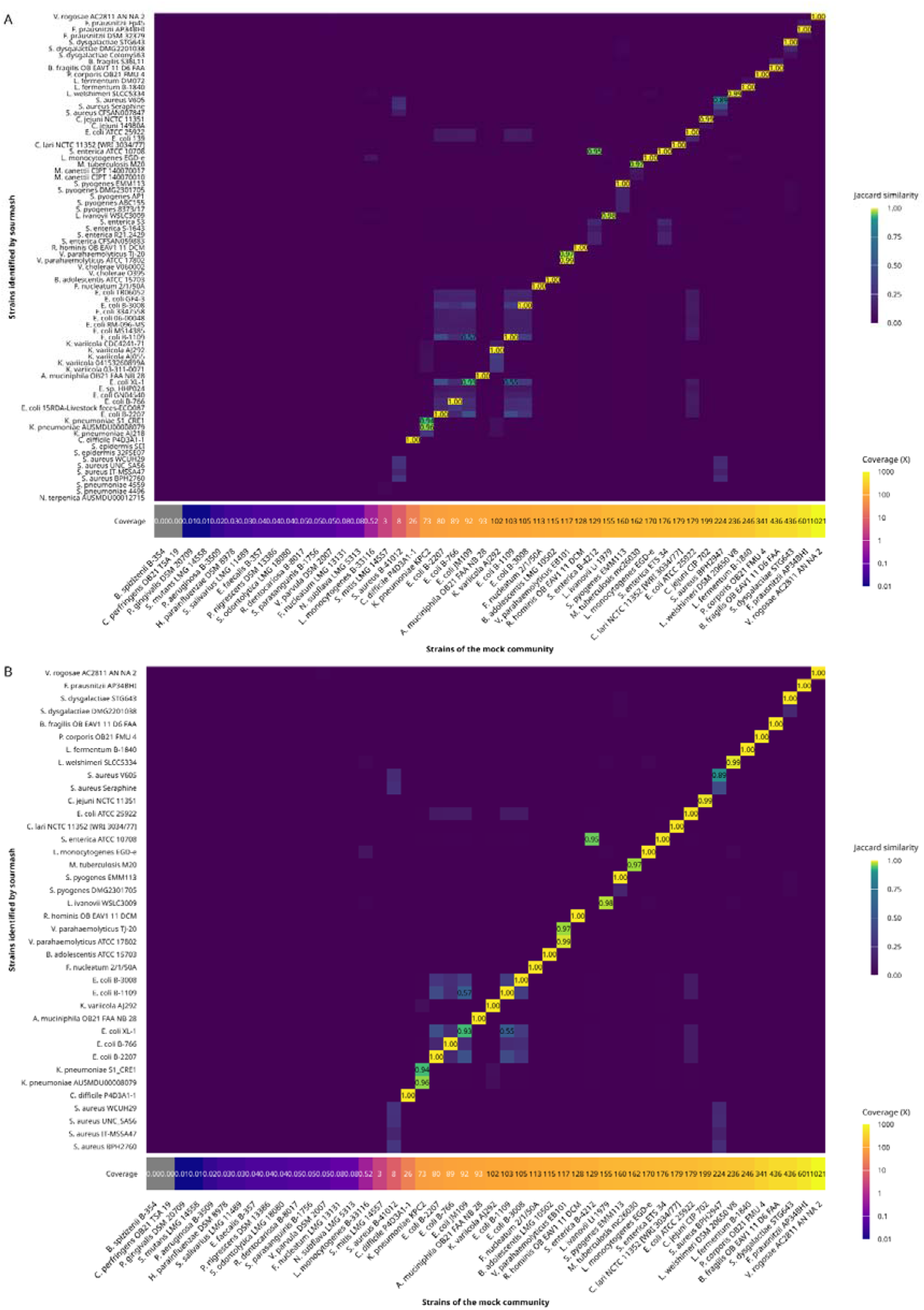
Gene content similarity heatmap between CONCAT mock community strains and strains identified by sourmash **(A)** using a minimum *reference fraction* of 0.5 **(B)** using a minimum *reference fraction* of 0.9.

As expected, results are slightly better with the three simpler communities HMW, ORAL, and GUT (Supplementary Figures 6, 7, and 8). Sourmash identified all expected strains in HMW and ORAL, and 16 out of the 18 expected strains in GUT, either as exact or closely related strains. The high identification rate in these communities may reflect their relatively low complexity and limited number of low-abundance strains. For matches that did not correspond to exact expected strain, these closely related strains have a gene content similarity of 0.93 with the expected strains of the mock communities. For the PANTAX community, 8 strains remain undetected, neither as exact nor as close matches. This happens even at coverages exceeding 2X, which is higher than the coverage at which the lowest abundant strains were identified (Supplementary Figure 9). Unidentified strains with a coverage over 2X could be explained by the presence, for each strain of the mock community, of another closely related strain of the same species. For example, *B. turicatae* BTE5EL is not identified at all, while the other strain of the same species in the mock, *B. turicatae* 91E135, is exactly identified and they have an ANI score of 99.8%. This highlights the limitation of strain-level identification: when genomes are highly similar, distinguishing between strains remains challenging. This limitation is not specific to sourmash but reflects a real difficulty of reference-based strain identification methods when several nearly identical genomes are present in the reference database.

Several strains reported by sourmash corresponded to false positives. The reference fraction given by sourmash for each strain identified was therefore used to filter the results. Among the tested thresholds, a value of 0.9 represented the optimal compromise between reducing false positives and preserving true positives (Supplementary Figure 5). For example, for the CONCAT mock community, using a minimal reference fraction of 0.5 resulted in 86 identified strains (Figure 3.A), whereas increasing this threshold to 0.9 reduced the number of identified strains to 38 (Figure 3.B). Even for the simpler mock communities, this threshold reduced the number of low-confidence genomes while preserving most expected strains. The total number of identified genomes decreased from 28 to 7 for HWM, from 14 to 12 for ORAL, and from 48 to 15 for GUT, substantially improving precision.

Overall, sourmash performance depends on sufficient genome coverage but, above this threshold, the majority of strains are recovered either as exact or closely related strains, sharing a large fraction of the expected gene content and supporting their use in downstream functional analyses.

### Sourmash accurately recovers community gene content using exact and close strains

In order to evaluate whether sourmash allows the recovery of the overall gene content of a community, the EggNOG orthologous groups of identified strains were compared to those of expected strains of mocks. For evaluation, recall, precision and F1-score have been calculated. These metrics were computed only for strains with a genome coverage above 3X, as those below this threshold were not reliably recovered in our framework. The three simplest mock communities, HMW, ORAL and GUT, have almost perfect results, with F1-scores over 0.99. This is consistent with their low complexity and shows that when strain diversity is limited, sourmash can recover the majority of the expected functional potential.

**Table 2:** Recall, precision and F1-score on gene content on the five mock communities. Metrics were calculated based on the gene content of mock strains with a coverage over 3X and strains identified by sourmash with a reference fraction over 0.9.

| **Mock communities** | **Recall** | **Precision** | **F1-score** |
|---|---|---|---|
| **HMW** | 0.996 | 0.997 | 0.997 |
| **ORAL** | 0.994 | 0.994 | 0.994 |
| **GUT** | 0.999 | 0.997 | 0.998 |
| **CONCAT** | 0.985 | 0.902 | 0.942 |
| **PANTAX** | 0.956 | 0.929 | 0.942 |

Even if they are more complex, CONCAT and PANTAX mock communities show high performance, with F1-scores of 0.942. The lower precision value for CONCAT (0.902) indicates that excess genes were predicted. They could come from the low abundant strains that were not included in the expected gene content. However, the recall value is over 0.98, indicating that the vast majority of expected genes of strains over 3X are recovered. It suggests that, despite some strains being absent from the detection, the functional potential of the community is preserved because closely related strains have highly similar gene content. The 3X threshold should be looked at as an empirical threshold coming from the sequencing conditions and mock communities used here rather than as an universal requirement for sourmash. Different sequencing depths of datasets may shift this limit. Even if the PANTAX mock community shows high recall and precision values despite its complexity, it should be interpreted with caution because reads are simulated. Moreover, it does not represent the complexity of real abundance variations found in a microbiota bacterial community. Mock communities provide controlled frameworks but rarely reproduce the complexity of real microbiomes, which are characterised by uneven abundances, unknown strains, and high diversity. Moreover, this study focuses on human gut and oral microbiomes, two ecosystems for which large reference genome databases are available. Consequently, sourmash performance is expected to be the highest in well-characterised environments. In ecosystems with a large part of undescribed diversity, such as soil microbiomes, the identification of biologically relevant strains would be more limited.

In summary, these results show that sourmash is able to recover the gene content of a microbial community either by identifying exact strains or closely related strains. Although orthologous groups provide only a functional approximation and don’t capture all differences in gene regulation or gene expression, they represent a relevant proxy for the functional potential of a microbial community. Thus, sourmash can support downstream analyses such as functional profiling or metabolic network reconstruction, even when the exact strain is not recovered.

### Accurate abundance estimation using sourmash identified genomes

To evaluate whether genomes identified by sourmash can be suitable references for abundance estimation, we compared the abundances of identified genomes and expected abundances of the mocks using the Pearson correlation. Ground truth abundances were estimated by mapping reads to the expected genomes of the mock communities, rather than using the theoretical mock composition, because sequencing biases can affect the initial abundances provided by the mock designs. Abundances of exact strains and strains identified by sourmash for each mock can be found in Supplementary Data 2.

To assess abundance estimation independently from strain identification accuracy, we compared the estimated and ground truth abundances only for strains that were exactly identified by sourmash. Across all five mock communities, abundance estimations were highly correlated with ground truth, with Pearson correlation coefficients ranging from 0.823 to 1 (Figure 4).

**Figure 4:**
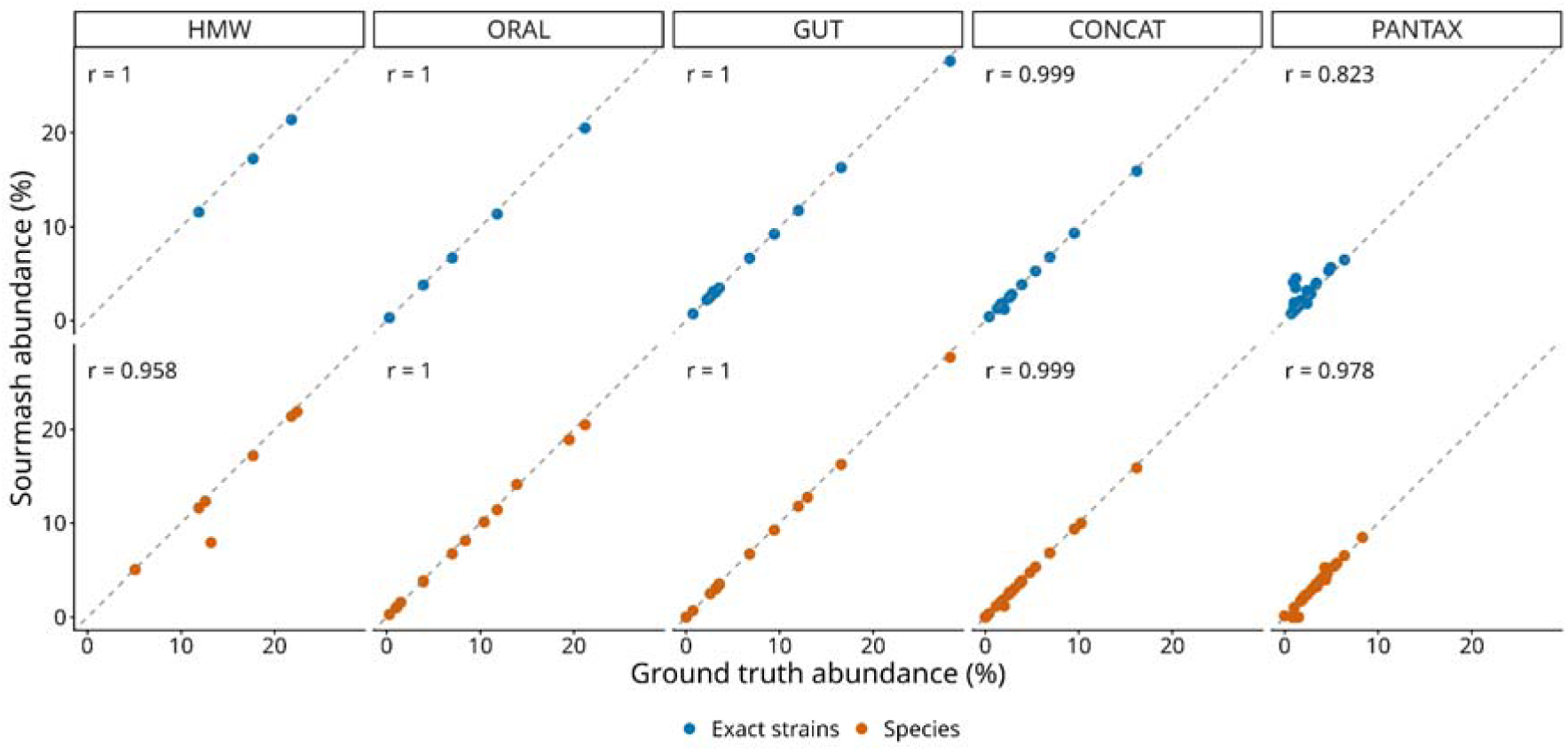
Correlation between ground truth and estimated abundances with sourmash identified genomes. Pearson correlations are shown for exact strain identifications and after aggregation of strains at the species level. High values of Pearson correlation coefficient (r) indicate an accurate abundance estimation.

However, as shown previously, sourmash often retrieves closely related strains rather than exact reference strains. To evaluate whether these strain-level differences affect abundance estimation, all identified strains abundances were aggregated at the species level. After this aggregation, correlations with the ground truth remained consistently high across all mock communities, ranging from 0.978 to 1 (Figure 4). This indicates that closely related strains have a limited impact on quantitative profiling.

Altogether, these results show that the main limitation of sourmash lies in discriminating between closely related strains rather than in abundance estimation. Genomes identified by sourmash can provide accurate quantitative information while also supporting strain-level functional characterisation through gene content similarity.

## Conclusions

In this study, we evaluated the ability of sourmash to identify bacteria at the strain level, to recover the gene content and to preserve abundance estimations of metagenomic samples, using five mock communities of increasing complexity with known compositions. We showed that nucleotide similarity alone is insufficient to capture the functional relatedness of two strains and that gene content similarity provides valuable complementary information. Above a sufficient genome coverage, sourmash identifies strains of a metagenomic sample, either exactly or with a closely related strain sharing a highly conserved gene content. This allows the recovery of the functional potential of the community. Furthermore, abundance estimation is consistent with the expected community composition, highlighting that the quantitative structure of the communities is preserved despite the identification of closely related strains.

This study was conducted exclusively on mock communities, which provide a known ground truth. However, mock communities do not fully reproduce the complexity of natural microbiomes, which contain a much larger diversity of organisms, uneven abundance distributions, and strains that may be absent from reference databases. Furthermore, this work focused only on gut and oral communities, two ecosystems for which large reference genome collections are available. Consequently, the performance reported in this study should be interpreted as an evaluation of sourmash under controlled conditions rather than as an estimate of its performance on real metagenomic samples. Future work should assess this workflow on real datasets of higher complexity to broaden these findings.

Overall, our results suggest that sourmash is well suited for strain-level metagenomic profiling in well-characterised ecosystems, such as the human gut microbiota, where representative reference genomes are available. Although distinguishing similar strains remains challenging, the identification of closely related strains with a conserved gene content and an accurate abundance estimation is a valuable alternative for downstream analyses such as functional profiling or metabolic reconstruction.

## Declarations

### Ethics approval and consent to participate

Not applicable.

### Consent for publication

Not applicable.

### Availability of data and material

The Nanopore sequencing reads of the three ZymoBIOMICS mock communities HMW, ORAL, and GUT, are publicly available in the European Nucleotide Archive (ENA) with the following accession numbers: ERR14294023, ERR14251413, ERR14251410, respectively. The CONCAT mock community was generated by merging the reads from these three previous ones and sequencing reads from 14 isolated strains, which are publicly available under the BioProject accession number PRJNA1087001. The simulated reads for the PANTAX mock community are publicly available on Zenodo: https://doi.org/10.5281/zenodo.15148106.

### Authors’ contributions

L.A. devised the study, ran the experiments, and drafted the manuscript. E.R. and R.V. co-supervised the study. All authors contributed regularly to practical decisions and solutions to explore. All authors edited the manuscript, read and agreed on the final version.

## Acknowledgments

We acknowledge the GenOuest bioinformatics core facility https://www.genouest.org/ for providing the computing infrastructure.

## References

1. Durazzi F, Sala C, Castellani G, Manfreda G, Remondini D, De Cesare A. Comparison between 16S rRNA and shotgun sequencing data for the taxonomic characterization of the gut microbiota. Sci Rep. 2021 Feb 4;11(1):3030. doi:10.1038/s41598-021-82726-y

2. Pinto Y, Bhatt AS. Sequencing-based analysis of microbiomes. Nat Rev Genet. 2024 Dec;25(12):829–45. doi:10.1038/s41576-024-00746-6

3. Jain C, Rodriguez-R LM, Phillippy AM, Konstantinidis KT, Aluru S. High throughput ANI analysis of 90K prokaryotic genomes reveals clear species boundaries. Nat Commun. 2018 Nov 30;9(1):5114. doi:10.1038/s41467-018-07641-9

4. Ghazi AR, Münch PC, Chen D, Jensen J, Huttenhower C. Strain Identification and Quantitative Analysis in Microbial Communities. J Mol Biol. 2022 Aug;434(15):167582. doi:10.1016/j.jmb.2022.167582

5. Orsi RH, Bakker HCD, Wiedmann M. Listeria monocytogenes lineages: Genomics, evolution, ecology, and phenotypic characteristics. Int J Med Microbiol. 2011 Feb;301(2):79–96. doi:10.1016/j.ijmm.2010.05.002

6. Fitz-Gibbon S, Tomida S, Chiu BH, Nguyen L, Du C, Liu M, et al. Propionibacterium acnes Strain Populations in the Human Skin Microbiome Associated with Acne. J Invest Dermatol. 2013 Sep;133(9):2152–60. doi:10.1038/jid.2013.21

7. 7. Zhang (•天•) T, Jiang (蒋冕) M, Li (李•洲) H, Gao (高云云) Y, Yousuf S, Yu (余•敏) K, et al. Computational Tools and Resources for Long-read Metagenomic Sequencing Using Nanopore and PacBio. Ning Z, editor. Genomics Proteomics Bioinformatics. 2025 Sep 22;23(4):qzaf075. doi:10.1093/gpbjnl/qzaf075

8. Mahmoud M, Agustinho DP, Sedlazeck FJ. A Hitchhiker’s Guide to long-read genomic analysis. Genome Res. 2025 Apr;35(4):545–58. doi:10.1101/gr.279975.124

9. Blanco-Miguez A, Beghini F, Cumbo F, McIver LJ, Thompson KN, Zolfo M, et al. Extending and improving metagenomic taxonomic profiling with uncharacterized species with MetaPhlAn 4 [Internet]. Bioinformatics; 2022 [cited 2026 Jul 17]. Available from: http://biorxiv.org/lookup/doi/10.1101/2022.08.22.504593 doi:10.1101/2022.08.22.504593

10. Olm MR, Crits-Christoph A, Bouma-Gregson K, Firek B, Morowitz MJ, Banfield JF. InStrain enables population genomic analysis from metagenomic data and rigorous detection of identical microbial strains [Internet]. Microbiology; 2020 [cited 2026 Jul 17]. Available from: http://biorxiv.org/lookup/doi/10.1101/2020.01.22.915579 doi:10.1101/2020.01.22.915579

11. Zheng A, Shaw J, Yu YW. Mora: abundance aware metagenomic read re-assignment for disentangling similar strains. BMC Bioinformatics. 2024 Apr 23;25(1):161. doi:10.1186/s12859-024-05768-9

12. Lipovac J, Angevin L, Križanović K. Using Mapping-Profiles to Refine Strain-Level Metagenomic Classification [Internet]. Bioinformatics; 2026 [cited 2026 Jul 22]. Available from: http://biorxiv.org/lookup/doi/10.64898/2026.05.18.725856 doi:10.64898/2026.05.18.725856

13. Lipovac J, Šikić M, Vicedomini R, Križanović K. MADRe: Strain-Level Metagenomic Classification Through Assembly-Driven Database Reduction [Internet]. Bioinformatics; 2025 [cited 2026 Jun 17]. Available from: http://biorxiv.org/lookup/doi/10.1101/2025.05.12.653324 doi:10.1101/2025.05.12.653324

14. Ondov BD, Treangen TJ, Melsted P, Mallonee AB, Bergman NH, Koren S, et al. Mash: fast genome and metagenome distance estimation using MinHash. Genome Biol. 2016 Dec;17(1):132. doi:10.1186/s13059-016-0997-x

15. Šošić M, Šikić M. Edlib: a C/C ++ library for fast, exact sequence alignment using edit distance. Hancock J, editor. Bioinformatics. 2017 May 1;33(9):1394–5. doi:10.1093/bioinformatics/btw753

16. Siekaniec G, Roux E, Lemane T, Guédon E, Nicolas J. Identification of isolated or mixed strains from long reads: a challenge met on Streptococcus thermophilus using a MinION sequencer. Microb Genomics. 2021 Nov 30;7(11). doi:10.1099/mgen.0.000654

17. Pierce NT, Irber L, Reiter T, Brooks P, Brown CT. Large-scale sequence comparisons with sourmash. F1000Research. 2019 Jul 4;8:1006. doi:10.12688/f1000research.19675.1

18. Rahman Hera M, Liu S, Wei W, S. Rodriguez J, Ma C, Koslicki D. Fast, lightweight, and accurate metagenomic functional profiling using FracMinHash sketches [Internet]. Bioinformatics; 2023 [cited 2026 Jun 17]. Available from: http://biorxiv.org/lookup/doi/10.1101/2023.11.06.565843 doi:10.1101/2023.11.06.565843

19. Shaw J, Marin MG, Li H. High-resolution metagenome assembly for modern long reads with myloasm. Nat Biotechnol. 2026 Mar 27. doi:10.1038/s41587-026-03053-z

20. Zhang W, Liu Y, Xu J, Chen E, Schönhuth A, Luo X. PanTax: Strain-level taxonomic classification of metagenomic data using pangenome graphs [Internet]. 2024 [cited 2025 Feb 18]. Available from: http://biorxiv.org/lookup/doi/10.1101/2024.11.15.623887 doi:10.1101/2024.11.15.623887

21. Fritz A, Hofmann P, Majda S, Dahms E, Dröge J, Fiedler J, et al. CAMISIM: simulating metagenomes and microbial communities. Microbiome. 2019 Dec;7(1):17. doi:10.1186/s40168-019-0633-6

22. Wood DE, Lu J, Langmead B. Improved metagenomic analysis with Kraken 2. Genome Biol. 2019 Nov 28;20(1):257. doi:10.1186/s13059-019-1891-0

23. Dong S, Kryczka A, Jin Y, Stumm M. RocksDB: Evolution of Development Priorities in a Key-value Store Serving Large-scale Applications. ACM Trans Storage. 2021 Nov 30;17(4):1–32. doi:10.1145/3483840

24. Cantalapiedra CP, Hernández-Plaza A, Letunic I, Bork P, Huerta-Cepas J. eggNOG-mapper v2: Functional Annotation, Orthology Assignments, and Domain Prediction at the Metagenomic Scale. Tamura K, editor. Mol Biol Evol. 2021 Dec 9;38(12):5825–9. doi:10.1093/molbev/msab293

25. Curry KD, Wang Q, Nute MG, Tyshaieva A, Reeves E, Soriano S, et al. Emu: species-level microbial community profiling of full-length 16S rRNA Oxford Nanopore sequencing data. Nat Methods. 2022 Jul;19(7):845–53. doi:10.1038/s41592-022-01520-4

26. Li H. Minimap2: pairwise alignment for nucleotide sequences. Birol I, editor. Bioinformatics. 2018 Sep 15;34(18):3094–100. doi:10.1093/bioinformatics/bty191

