## Supplementary Material for "Sourmash accurately profiles strains from long-read metagenomes preserving gene content"

Supplementary Figure 1: Characterisation of HMW mock community composition. **(A)** ANI scores heatmap of the strains of the mock community HMW. **(B)** Gene content similarity heatmap of gene content between HMW mock community strains. Below an ANI score of 70%, the exact score was not calculated by fastANI.
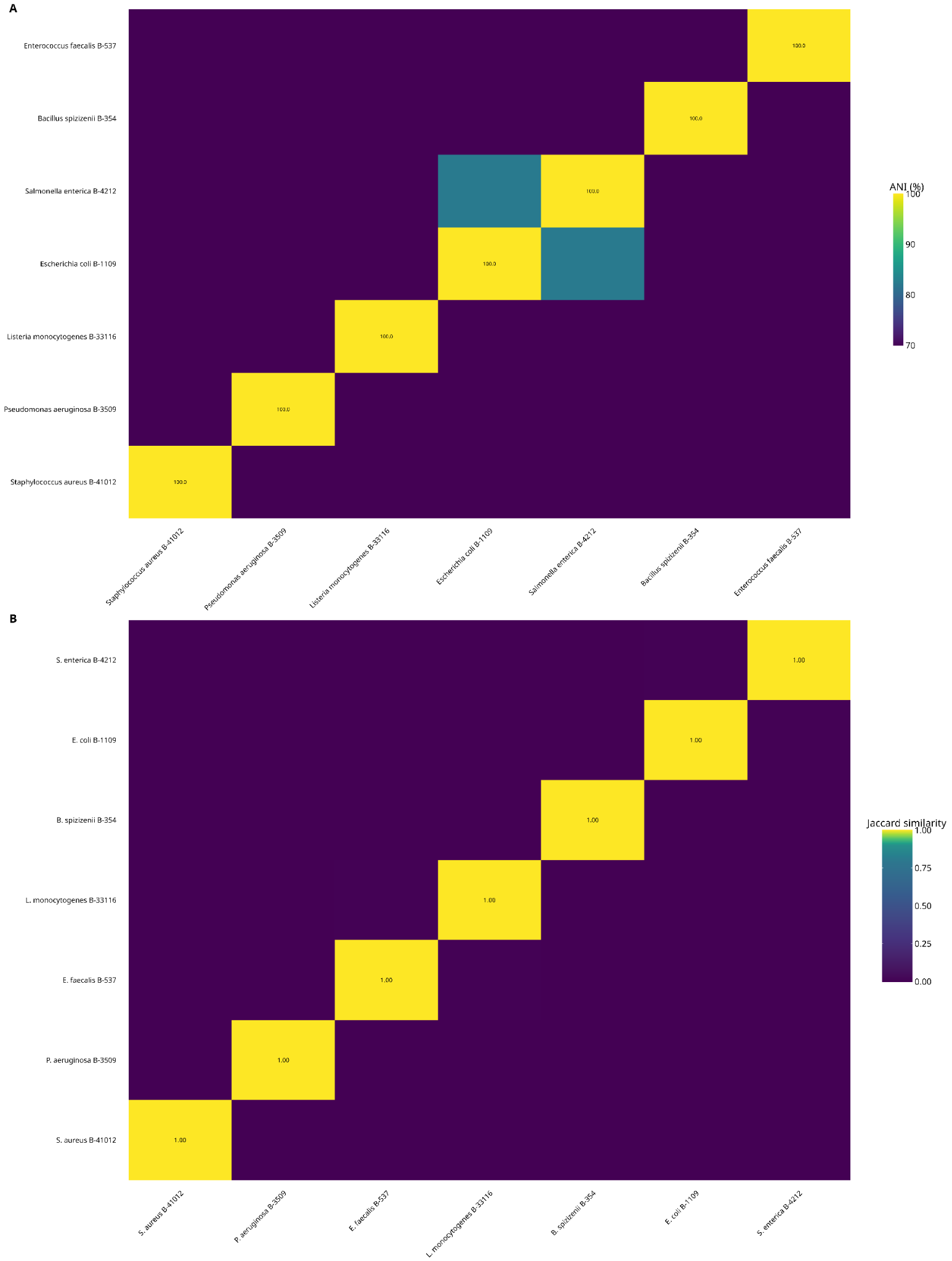

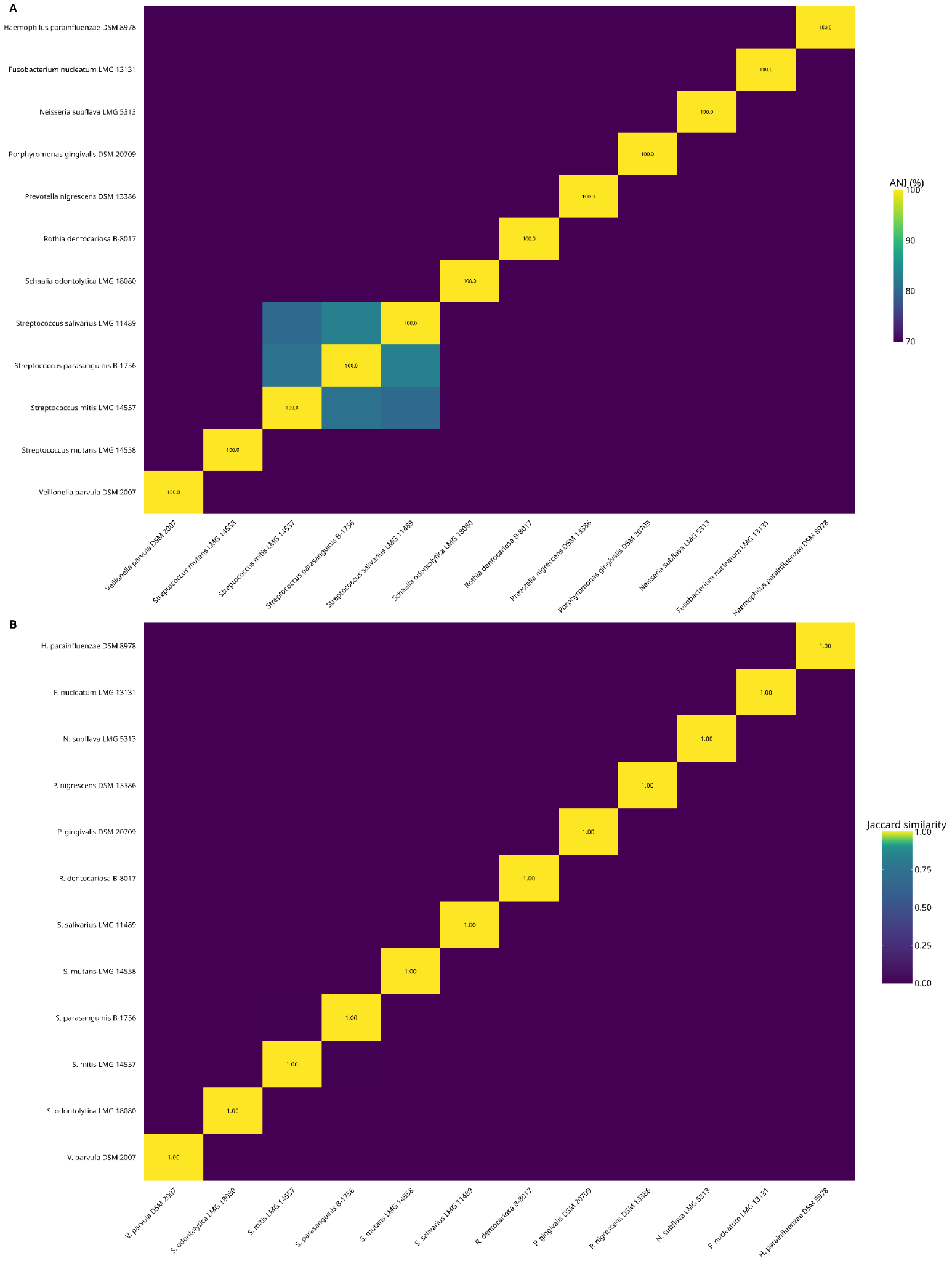
Supplementary Figure 2: Characterisation of ORAL mock community composition. **(A)** ANI scores heatmap of the strains of the mock community ORAL. **(B)** Gene content similarity heatmap of gene content between ORAL mock community strains. Below an ANI score of 70%, the exact score was not calculated by fastANI.

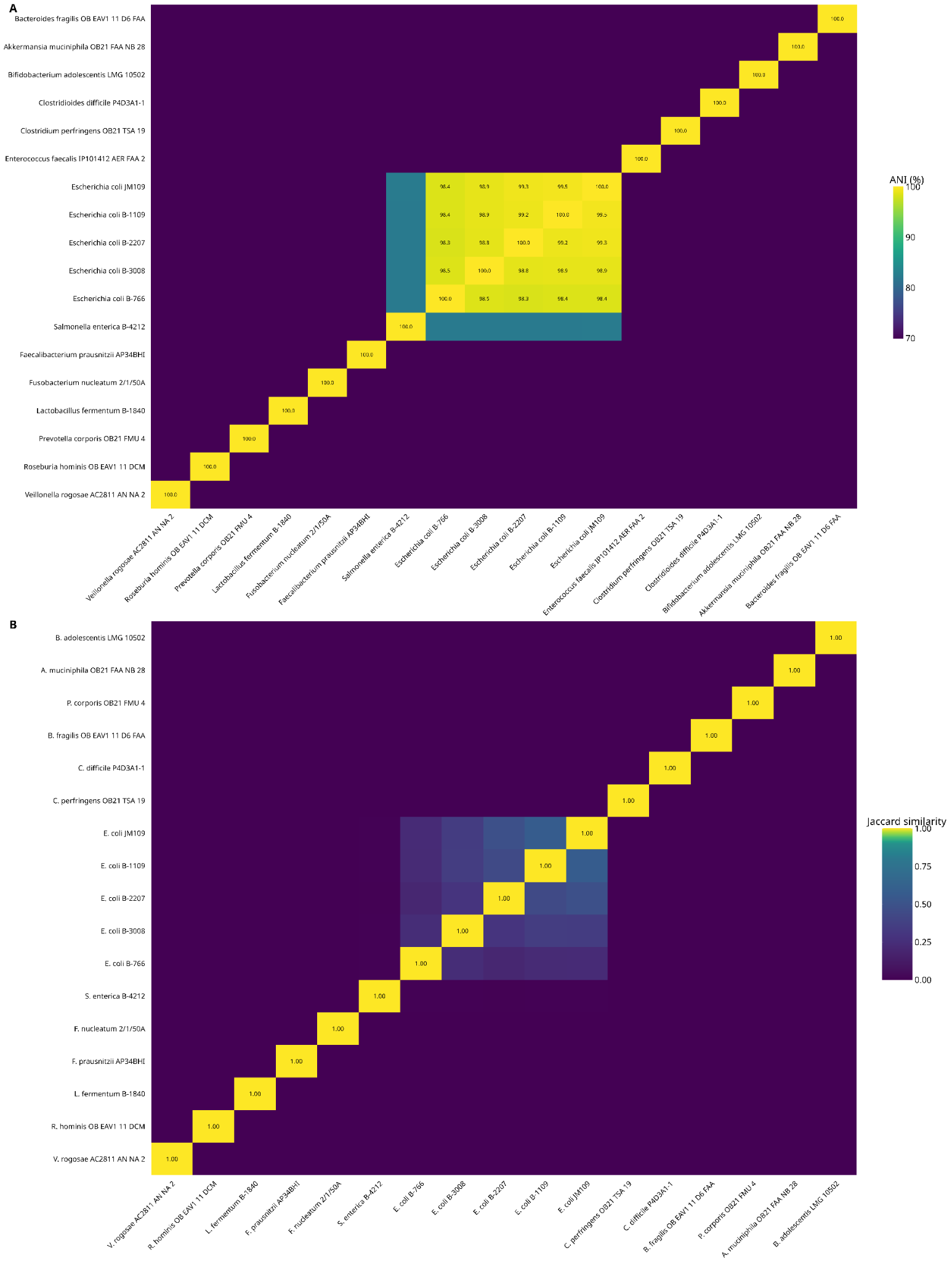
Supplementary Figure 3: Characterisation of GUT mock community composition. **(A)** ANI scores heatmap of the strains of the mock community GUT. **(B)** Gene content similarity heatmap of gene content between GUT mock community strains. Below an ANI score of 70%, the exact score was not calculated by fastANI.

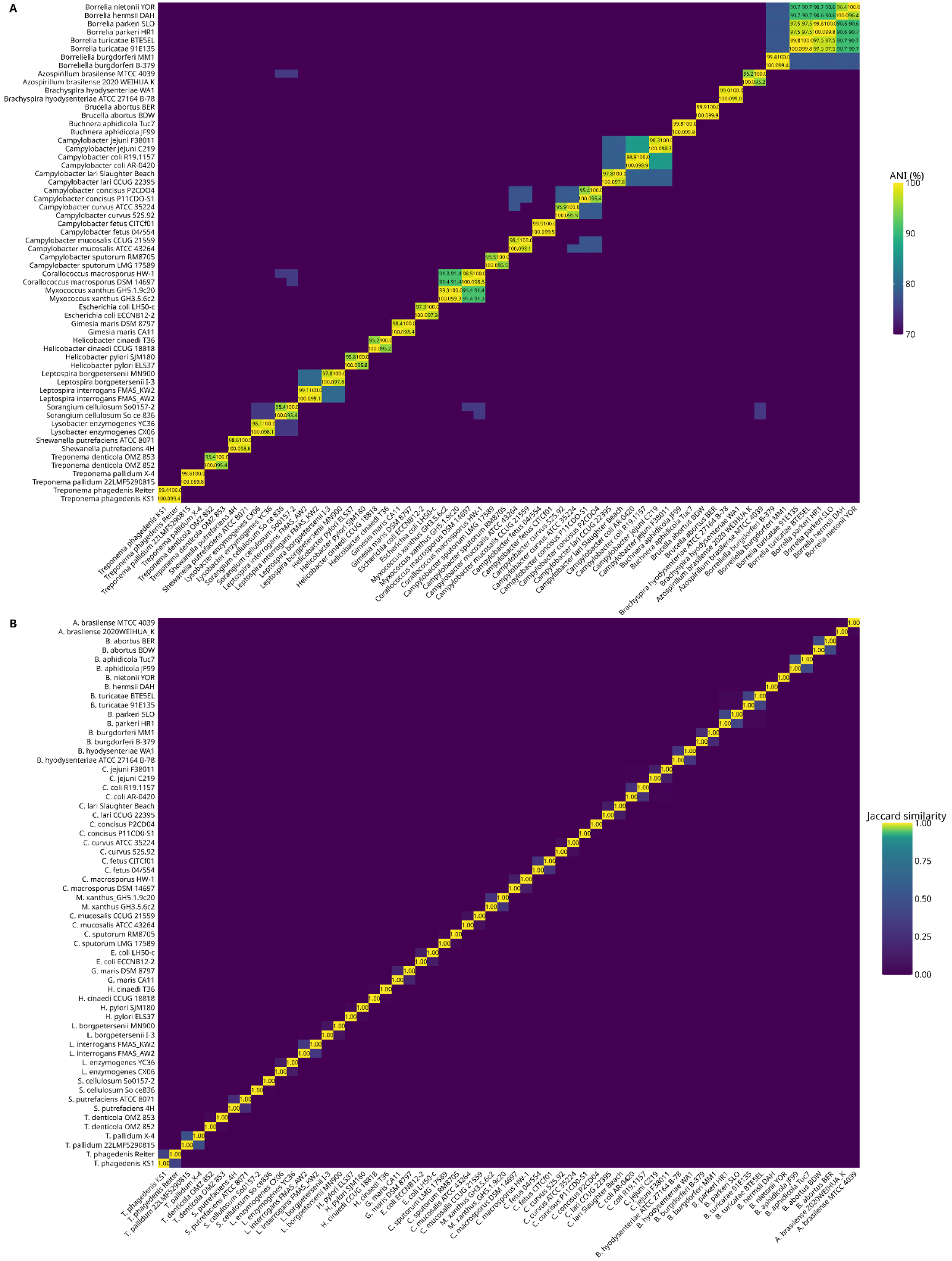

Supplementary Figure 4: Characterisation of PANTAX mock community composition. **(A)** ANI scores heatmap of the strains of the mock community PANTAX. **(B)** Gene content similarity heatmap of gene content between PANTAX mock community strains. Strains of the same species have high ANI scores, over 90%. Below an ANI score of 70%, the exact score was not calculated by fastANI.

Supplementary Figure 5: Evolution of F1-score across reference fraction values, databases and input sequences types for the five mock communities.
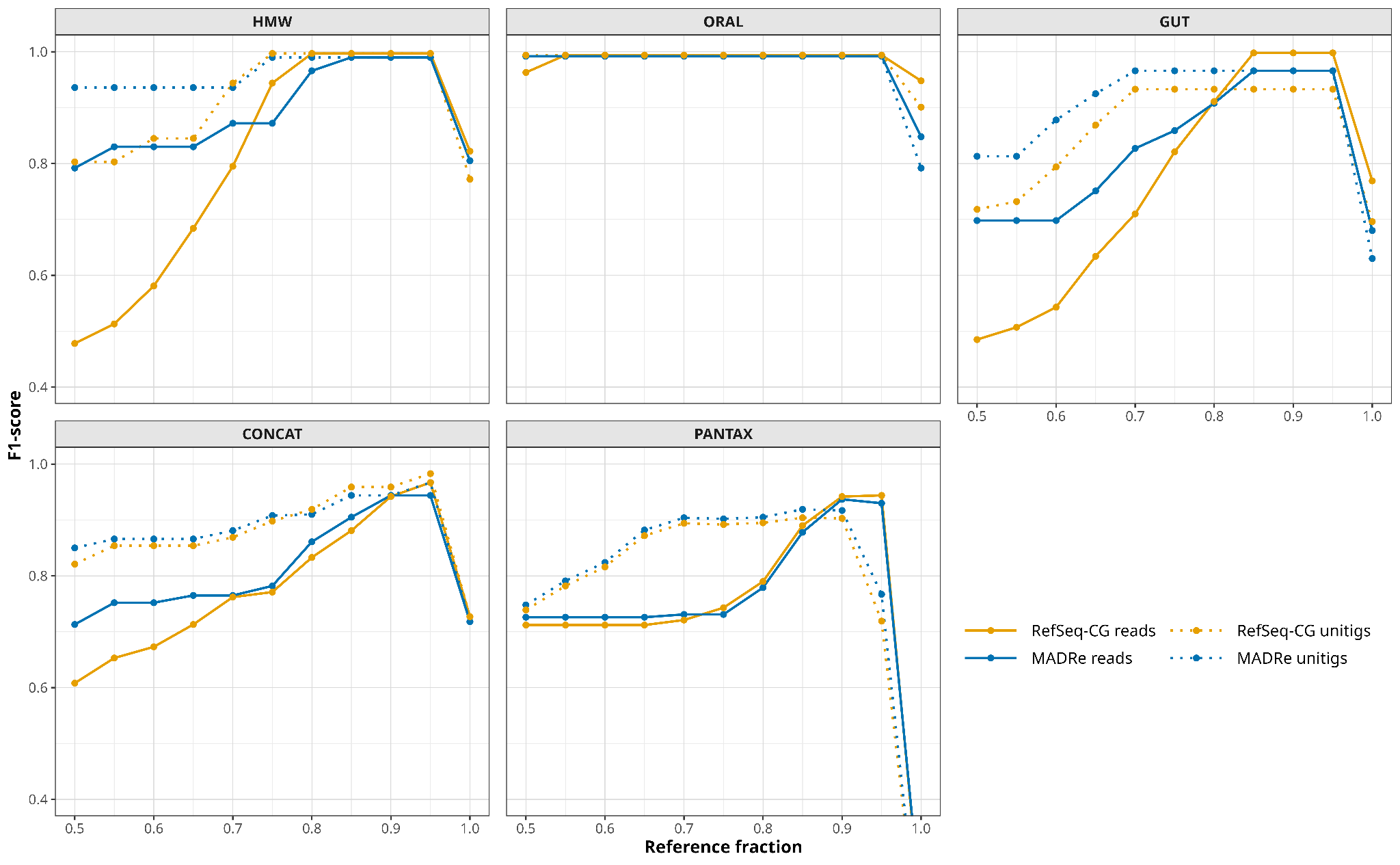

Supplementary Figure 6: Gene content similarity heatmap between HMW mock community strains and strains identified by sourmash **(A)** using a minimum reference fraction of 0.5 **(B)** using a minimum reference fraction of 0.9.
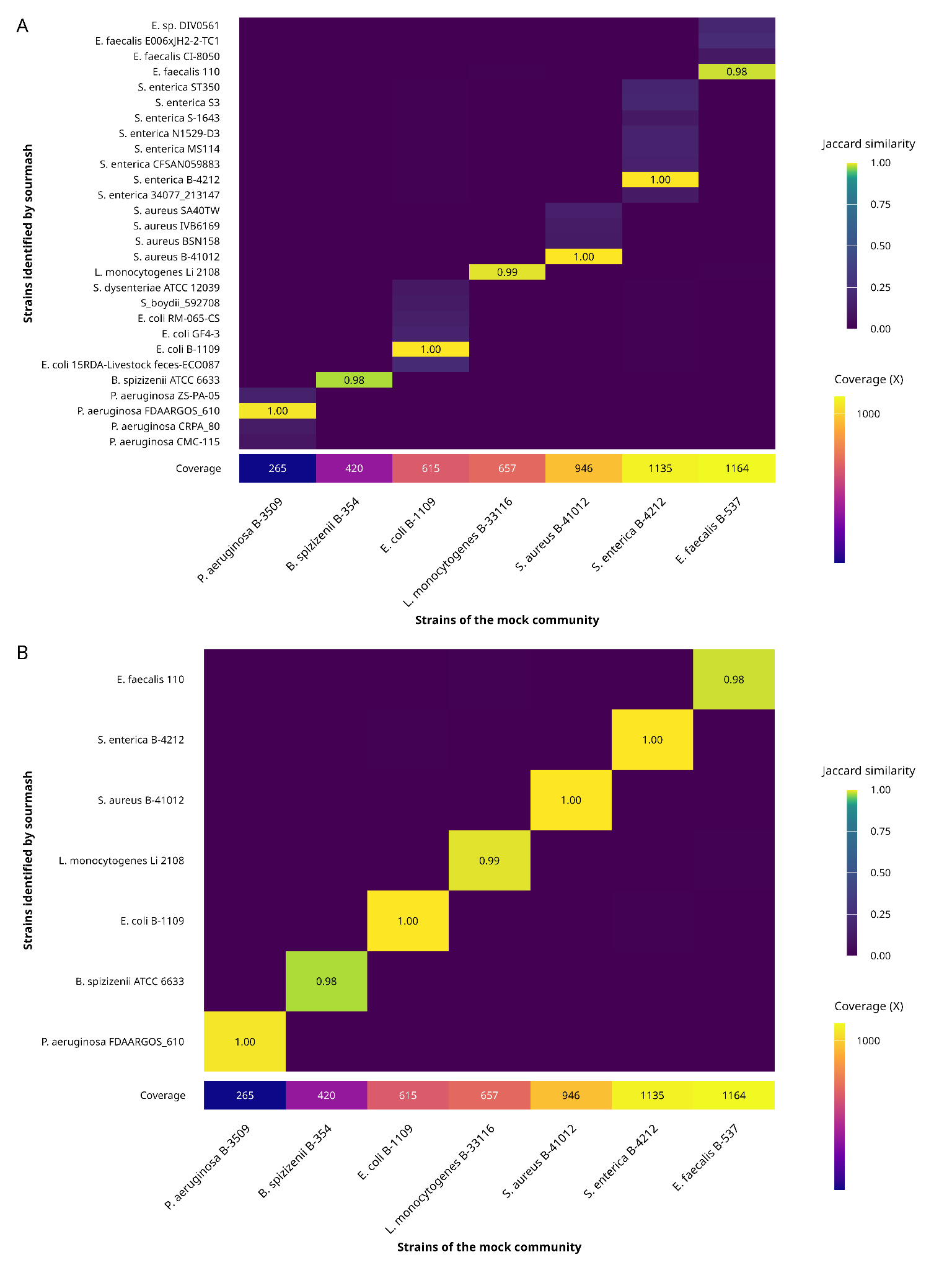

Supplementary Figure 7: Gene content similarity heatmap between ORAL mock community strains and strains identified by sourmash **(A)** using a minimum reference fraction of 0.5 **(B)** using a minimum reference fraction of 0.9.
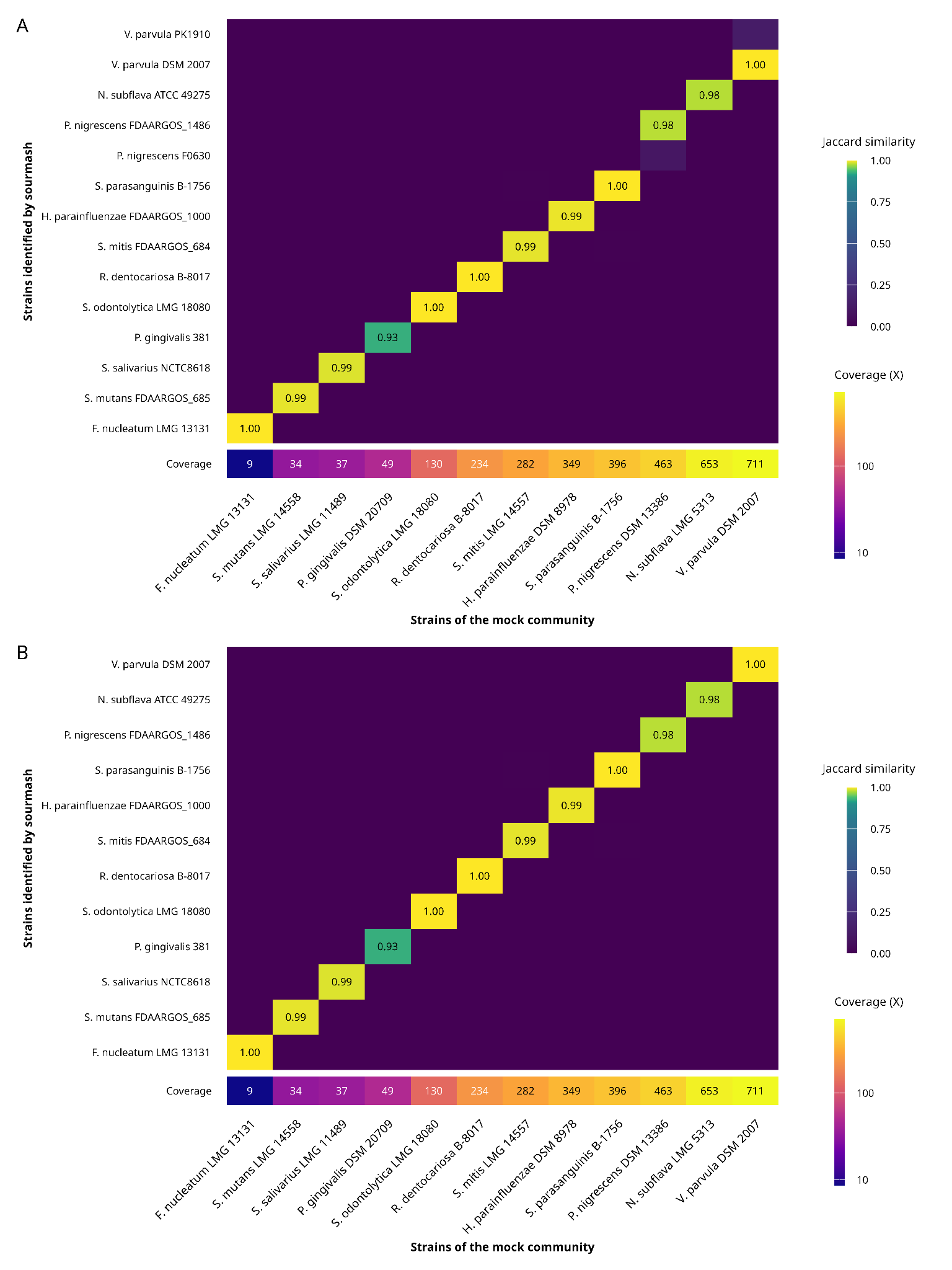

Supplementary Figure 8: Gene content similarity heatmap between GUT mock community strains and strains identified by sourmash **(A)** using a minimum reference fraction of 0.5 **(B)** using a minimum reference fraction of 0.9.
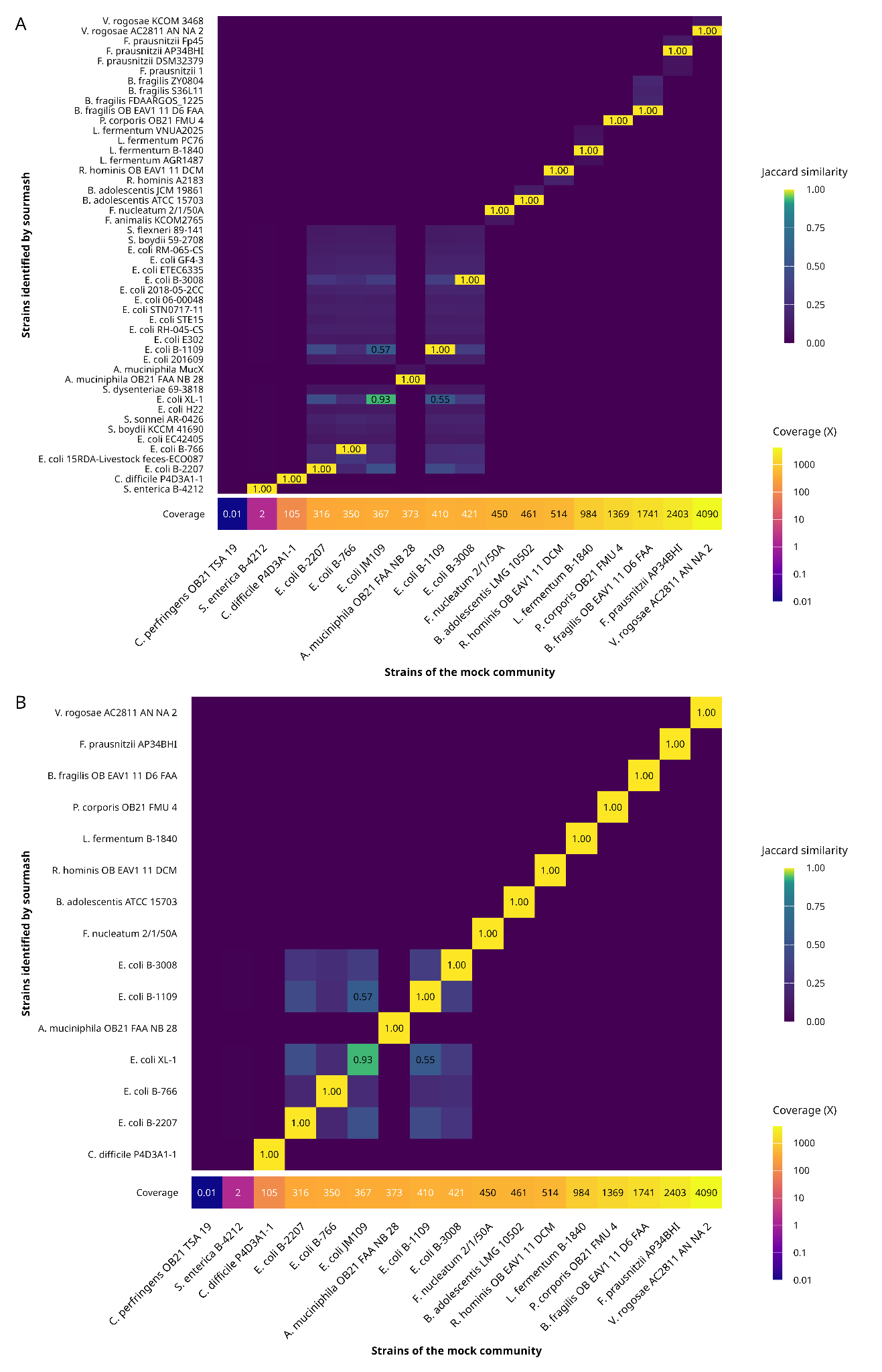

Supplementary Figure 9: Gene content similarity heatmap between PANTAX mock community strains and strains identified by sourmash **(A)** using a minimum reference fraction of 0.5 **(B)** using a minimum reference fraction of 0.9.
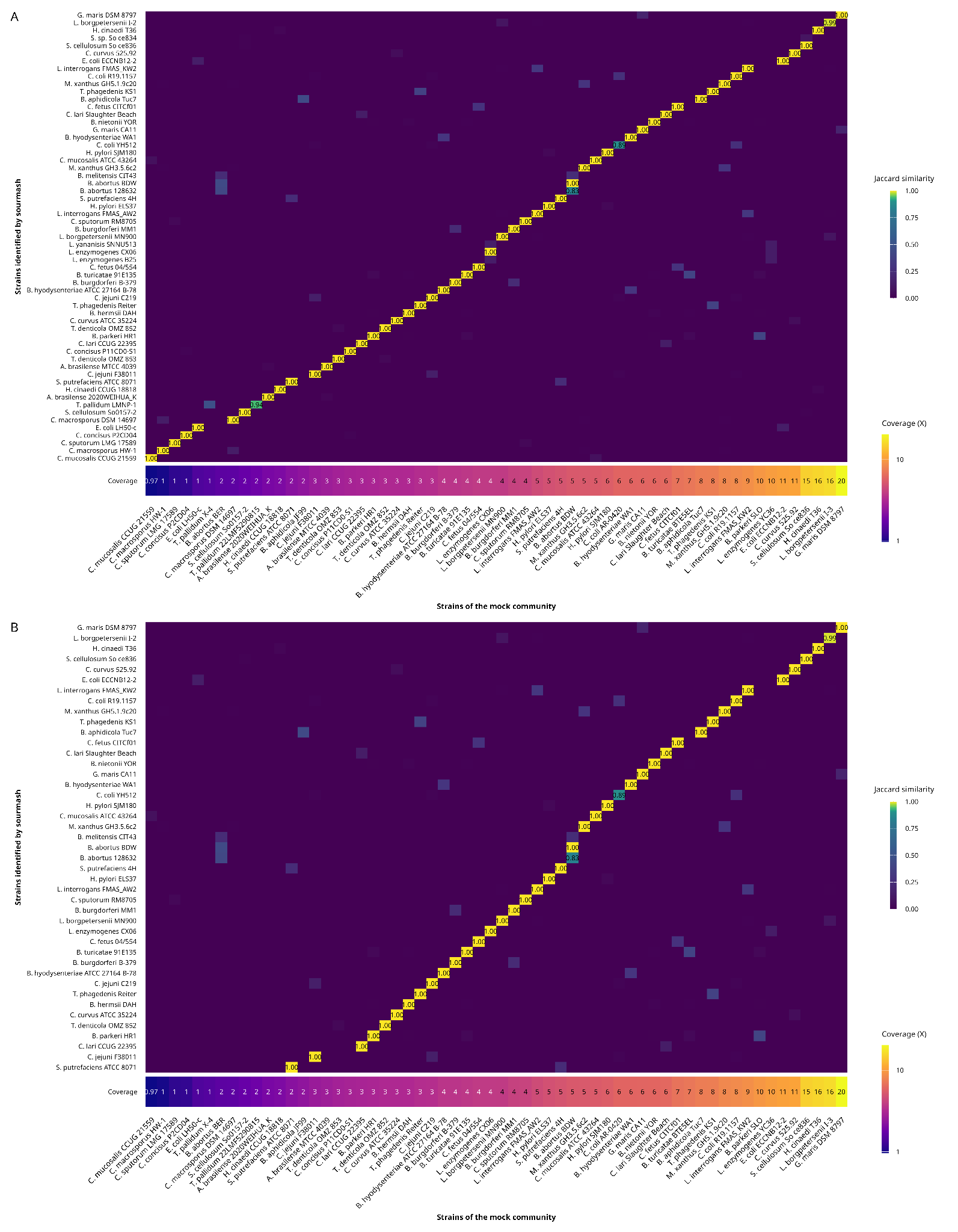
